# Mechanical strain of the intestinal epithelium directs absorptive lineage maturation

**DOI:** 10.64898/2026.07.08.737211

**Authors:** Ronja M. Houtekamer, Björn van Sambeek, Karen B. van den Anker, Marjolein J. Vliem, Rutger N.U. Kok, Mirjam C. van der Net, Maria Rodriguez-Colman, Alexander van Oudenaarden, Martijn Gloerich

## Abstract

The intestinal epithelium is continuously subjected to a variety of mechanical forces, including extrinsic peristaltic contractions and intrinsic tensile forces generated by epithelial cell migration. Yet, how these mechanical cues influence the cellular processes underlying intestinal homeostasis remains poorly understood. In this study, we examine the impact of mechanical forces on intestinal cell dynamics by applying controlled external stretch to intestinal organoids, combined with high-throughput single-cell transcriptomic profiling. Our analyses reveal that prolonged cyclic mechanical strain alters the composition of differentiated intestinal cell populations. Specifically, we identify a strain-induced shift in the absorptive lineage towards a less mature state, with expansion of the population of early-stage enterocytes at the crypt–villus interface. This shift is associated with downregulation of transcriptional programs controlling enterocyte maturation within absorptive precursor populations. Our findings indicate that mechanical strain directs the maturation of the intestinal absorptive lineage, and highlight a role for mechanical forces in shaping intestinal epithelial composition and function.

## Introduction

The homeostatic regulation of epithelial tissues relies on the capacity of cells to dynamically adapt their behavior in response to biochemical and physical signals from their environment. Mechanical forces are key regulators of cellular processes fundamental to epithelial homeostasis, including proliferation, fate specification, migration, and cell loss (Iskratsch et al., 2014; Swaminathan & Gloerich, 2021). These forces originate both within the tissue, for instance, when neighboring cells exert tension on each other, and from a variety of extrinsic sources. Cells can respond to mechanical cues through mechanosensitive proteins and complexes, which transduce forces into intracellular biochemical signals that instruct cellular behavior (Di et al., 2023; Swaminathan & Gloerich, 2021). Although substantial progress has been made in understanding cellular mechanosensing, how mechanical forces contribute to the regulation of epithelial homeostasis at the tissue level remains incompletely understood.

The adult small intestine is an archetypal model to study regulation of epithelial homeostasis, due to its coordinated and dynamic regulation of epithelial self-renewal, composition, and function (Choo et al., 2022; Gehart & Clevers, 2019). Intestinal epithelial cells are spatially organized into crypt and villus compartments that orchestrate local generation, differentiation, and extrusion of cells (Beumer & Clevers, 2020). Absorptive enterocytes along the villus ensure efficient nutrient uptake, while secretory cells (i.e., goblet, paneth, enteroendocrine, and tuft cells) support digestive and protective functions. These differentiated cell populations are continuously replenished by proliferative stem and progenitor cells in the crypt. As newly generated differentiated cells exit the crypt, they progressively mature and acquire specialized functions during their migration toward the villus tip, where they are eventually shed (Harnik et al., 2024; Moor et al., 2018). While the contributions of growth factor signaling (e.g., Wnt, Bmp, and Notch) to these intestinal processes is well established (Gehart & Clevers, 2019; Spit et al., 2018), how mechanical cues are integrated with these pathways to shape intestinal homeostasis remains largely unresolved.

The intestinal epithelium experiences a variety of mechanical forces (Houtekamer et al., 2022; Pérez-González et al., 2022). Extrinsic forces arise from peristaltic muscle contractions, villus motility, and luminal shear during food digestion (Mercado-Perez & Beyder, 2022). In parallel, intrinsic epithelial contractility generates tension within the crypt and villus compartments (Krueger et al., 2025; Pérez-González et al., 2021; Yang et al., 2021), and cell migration toward the villus tip establishes gradients of intercellular tension along the villus (Krndija et al., 2019). Forces can directly modulate the function of differentiated intestinal cell types, for instance, stimulating hormone and mucus production by enteroendocrine and goblet cells (Alcaino et al., 2018; Fang et al., 2023; Treichel et al., 2021; Xu et al., 2021). In the *Drosophila* midgut, digestion-associated forces have been shown to promote stem cell proliferation and maturation of enteroendocrine progenitors (He et al., 2018; Q. Li et al., 2018). In the mammalian intestine, modulation of forces and downstream mechanosensing processes uncovered a role for mechanics in regulating crypt cell proliferation and stem cell self-renewal, crypt fission, and villus epithelial cell extrusion (Baghdadi et al., 2024; Fernández-Sánchez et al., 2015; Hinnant et al., 2024; Krueger et al., 2025; Y. Li et al., 2020; Meng et al., 2022; Poling et al., 2018; Spencer et al., 2006; Tallapragada et al., 2021). Despite these advances, a comprehensive understanding of how mechanical cues regulate epithelial lineage composition across the full spectrum of intestinal cell types is lacking.

Here, we investigate how mechanical forces influence intestinal epithelial cell type composition by applying controlled extrinsic mechanical stretch to murine small intestinal organoids, which recapitulate the composition and compartmentalization of the native intestine. By combining this with single-cell transcriptome analyses, we found that prolonged cyclic mechanical strain alters the cellular composition of the intestinal epithelium. Mechanical strain reduces the abundance of enterocytes while increasing the number of non-specialized proliferative cells. We specifically demonstrate that mechanical strain shifts the absorptive lineage toward a less mature state. Taken together, our data identifies mechanical strain as a key regulator of intestinal epithelial lineage maturation, revealing a force-dependent mechanism that shapes intestinal composition and function.

## Results

### Stretch application to intestinal organoids induces mechanical strain along crypt-villus compartments

To investigate how intestinal epithelial cells respond to forces, we developed an approach to apply external stretch to intestinal organoids **(figure 1A, S1A)**. Crypt segments derived from the murine small intestine were embedded in basement membrane extract (BME) that was covalently coupled to the surface of a flexible polydimethylsiloxane (PDMS) membrane. After organoids developed into three-dimensional budded structures, mechanical strain was applied through motor-controlled uniaxial extension of the PDMS membrane. Transmission of tensile forces through the matrix resulted in deformation of the organoids **(figure 1B)**.

**Figure 1.**
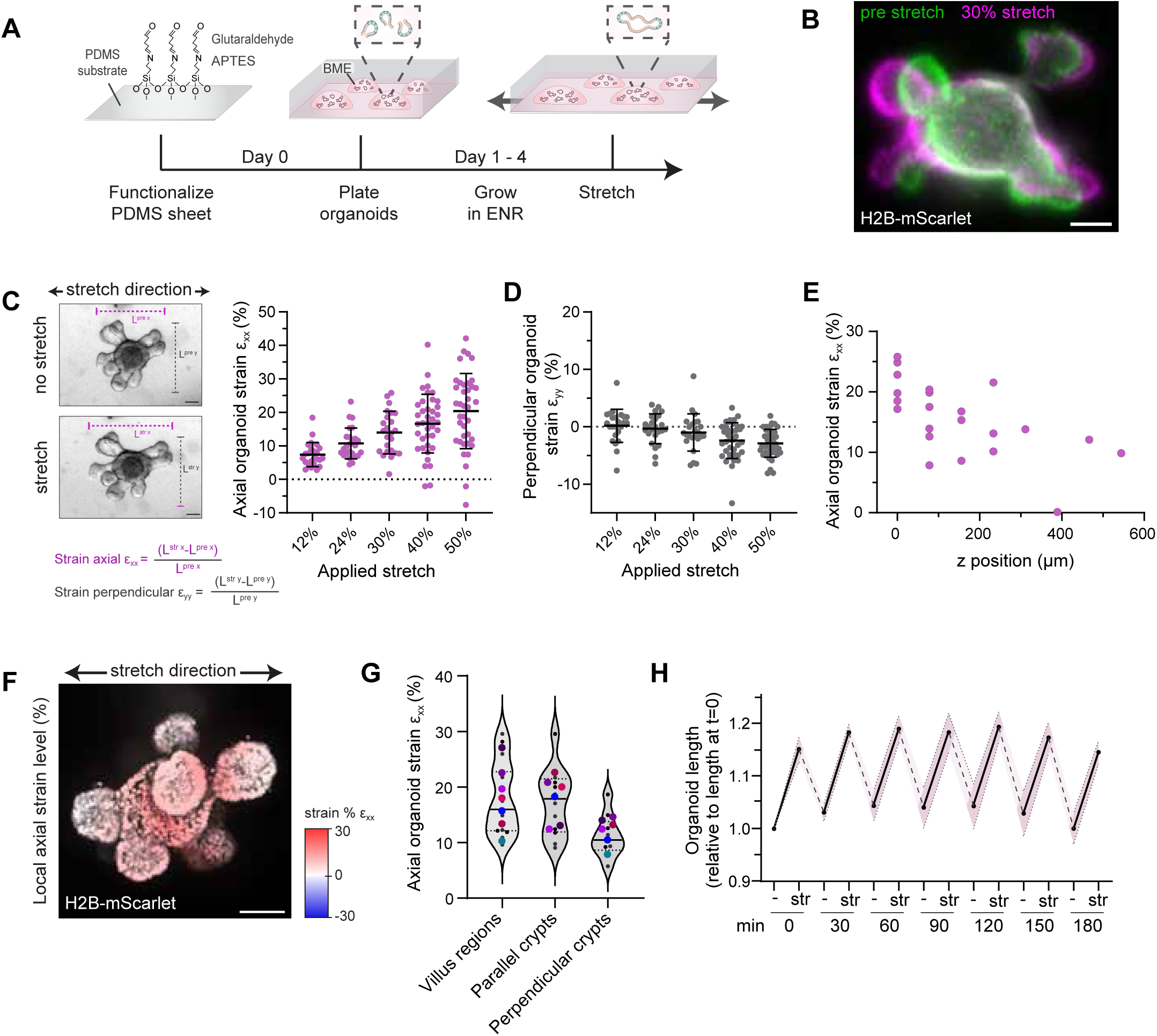
Characterization of intestinal organoid strain in response to mechanical stretch. **(A)** Experimental workflow of mechanical stretch application to mouse small intestinal organoids. Organoids embedded in basement membrane extract (BME) domes covalently coupled to functionalized PDMS membranes, and following 4 day culture in ENR differentiation medium and formation of budded structures, were subjected to static or cyclic uniaxial stretch. ENR: EGF/Noggin/R-spondin. **(B)** Representative overlay image of mouse small intestinal organoid overexpressing H2B-mScarlet-I before (green) and during 30% static stretch application (magenta). Scale bar: 50 μm. **(C)** Left: brightfield images of organoid before and during 30% static stretching. Scale bar: 50 μm. Right: corresponding quantification of whole-organoid axial strain (εxx, parallel to the stretch axis) with increasing magnitudes of stretch that were sequentially applied to the same organoids. Data represent mean with SD of n=24 (12, 24 and 30%) or n=39 (40 and 50%) organoids from 2-3 BME domes. **(D)** Quantification of strain perpendicular to the stretch axis (εyy) of organoids shown in figure 1C. **(E)** Axial strain percentages of budded organoids as a function of their distance to the PDMS (z = 0 μm represents the position of BME-embedded organoids closest to the PDMS) during 30% static stretch. Data represent individual budded organoids (n = 22) across 8 z slices from 4 BME domes. **(F)** Maximum intensity z-projection showing color-coded axial strain percentage of a representative intestinal organoid following 30% static stretch application. Scale bar: 50 μm. **(G)** Quantification of axial strain of villus regions, crypts parallel to the stretch axis and crypts perpendicular to the stretch axis (n=7 organoids). Colored data points represent means of crypt or villus regions per organoid**. (H)** Axial strain over time during cyclic stretching (0.1 Hz). Data represent mean relative lengths of n=5 individual organoids from 3 BME domes.

Quantitative analysis of whole-organoid deformation revealed that organoids experienced strain along the stretch axis, and the extent of this axial strain gradually increased with the applied stretch magnitude **(figure 1C)**. At stretch levels exceeding 30%, organoids exhibited 1-3% compression in the direction perpendicular to the stretch axis, but this was neglectable at lower stretch levels **(figure 1D)**. Despite a global correlation between applied stretch and organoid deformation, individual organoids displayed variability in their strain levels **(figure 1C)**. While this variation may in part be due to differences in organoid size and shape **(figure S1B)**, the proximity of organoids to the PDMS substrate was the predominant factor determining strain levels **(figure 1E)**. Organoids located closest to the substrate experienced highest axial strain, with strain levels decreasing progressively at larger distances **(figure 1E)**. Importantly, most organoids reside near the base of the matrix and thus experience robust and consistent axial strain that closely matches the applied stretch **(figure S1C)**.

To spatially resolve mechanical strain patterns within individual organoids, we performed live imaging of organoids expressing H2B-mScarlet-I during stretch application and measured local strain across organoid compartments. This revealed that both crypt and villus regions experienced substantial amounts of axial strain, although the exact strain levels within each crypt depended on its orientation relative to the stretch axis **(figure 1F-G)**. Thus, despite inter-compartment heterogeneity in strain magnitude, the entire intestinal organoid epithelium is subjected to mechanical strain.

Finally, we examined organoid strain dynamics during prolonged cyclic stretch application to assess whether organoids continued to experience stretch over time without adaptations. When applying cyclic stretch at a frequency of 0.1 Hz, organoids consistently experienced similar strain levels over several hours **(figure 1H, S1D)**. Thus, our cyclic stretching approach provides a robust platform to probe long-term downstream responses of organoids to mechanical strain.

### Prolonged cyclic mechanical strain triggers a reduction of enterocytes

Having characterized the stretch-induced mechanical strain patterns of intestinal organoids, we next sought to elucidate downstream mechanoresponses. To examine how mechanical strain influences intestinal cell type composition and identify potential underlying transcriptional responses, we subjected organoids to 15% or 30% cyclic stretch and performed single-cell RNA sequencing after 24 and 48 hours **(figure 2A)**. Two biological replicates were included to account for variability in stretch responses. Stretched organoids did not exhibit detectable changes in morphology and structural integrity compared to non-stretched organoids **(figure 2B)**. Moreover, no increase in apoptotic events was observed **(figure S2A)**. After quality control and filtering of the sequencing data, we obtained transcriptomes for 105.865 cells (approximately 9.000 cells per sample, with an average of 1916 genes and 3445 UMIs detected per cell). Unsupervised clustering of all cells combined identified 10 distinct clusters **(figure 2C, S2D)**. Annotation based on established mouse intestinal cell-type signatures **(Haber et al., 2017)** confirmed the presence of all expected adult intestinal cell types across each condition and at both timepoints **(figure 2C-D, S2D-F)**, enabling quantitative analysis of strain-induced changes across cell types.

**Figure 2.**
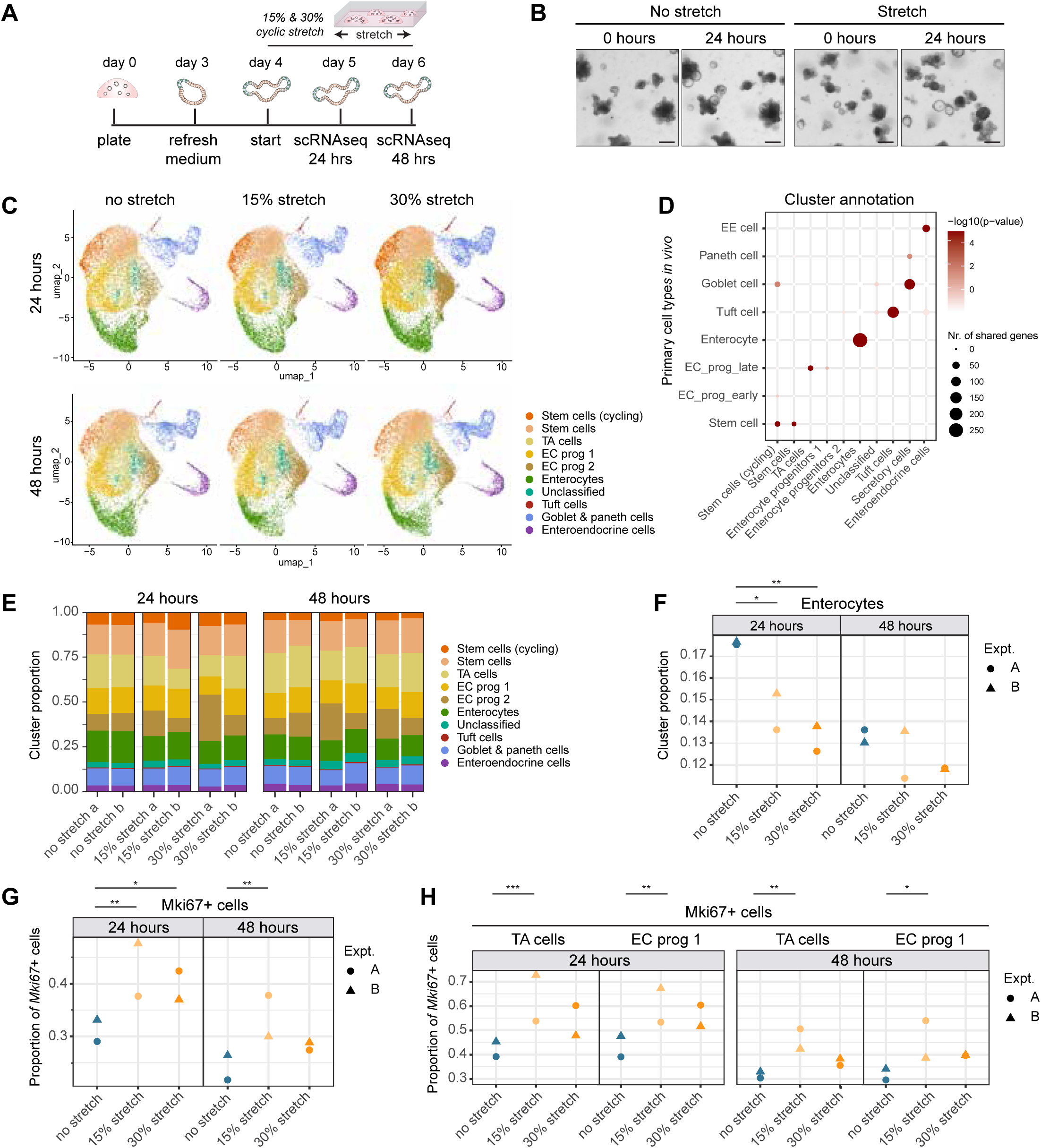
scRNAseq reveals that mechanical strain decreases the number of enterocytes in intestinal organoids. **(A)** Experimental workflow for single-cell RNA sequencing of mouse small intestinal organoids exposed to cyclic uniaxial mechanical stretch (0.1 Hz) or unstretched controls. Two independent biological replicates (A and B) were processed and sequenced in parallel. **(B)** Representative brightfield images of organoids cultured with or without 30% cyclic uniaxial stretch for 24 hours. Scale bar: 200 μm. **(C)** UMAP projection of single cell transcriptomic data from organoids from indicated conditions. Cluster identities were assigned based on transcriptional similarities to murine intestinal cell type signatures (Haber et al., 2017). Data represent 105.865 total single cells from two biological replicates, with ± 9.000 cells per sample**. (D)** Dot plot comparing organoid clusters, based on data from stretched and non-stretched organoids, to in vivo murine intestinal cell type clusters (Haber et al., 2017), showing number of shared genes and significance thereof, calculated using a binomial test. **(E)** Stacked bar plots showing the relative proportions of each annotated cell type cluster per sample across all conditions (non-stretched, 15%, 30% stretch, at 24 and 48 hours). Independent biological replicates (A and B) are plotted separately. **(F)** Proportions of enterocytes per sample across all conditions and timepoints. Replicate experiments are indicated by symbol shape. Statistics: Propeller method (linear model with variance stabilization and empirical Bayes moderation), as previously described (Phipson et al., 2022). * p < 0.05; ** p < 0.01**. (G)** Proportions of cells with expression of Mki67 for each condition and timepoint. Replicate experiments are indicated by symbol shape. Statistics: Binomial generalized linear mixed-effects model with Bonferroni-corrected p-values. * p < 0.05; ** p < 0.01; *** p < 0.001. (H) Proportion of Mki67-expressing cells within the indicated cell type clusters across all conditions and timepoints. Replicate experiments are indicated by symbol shape. Statistics: Binomial generalized linear mixed-effects model with Bonferroni-corrected p-values. * p < 0.05; ** p < 0.01; *** p < 0.001

In absence of applied stretch, cell type proportions were similar between replicates and showed only minor variability between both timepoints **(figure 2E)**. In contrast, mechanical strain resulted in two major changes in cellular composition. Most prominently, organoids subjected to either 15% or 30% stretch showed a significant reduction in the number of enterocytes **(figure 2F, S2G)**. This decrease was observed at both 24 and 48 hours, but was more pronounced and consistent between experiments at 24 hours **(figure 2F, S2G)**. Second, mechanical strain induced an increased prevalence of proliferating cells as marked by expression of *Mki67* **(figure 2G)**, and an increase of other proliferation-associated genes **(S3A).** This strain-induced proliferative response occurred across multiple cell types, but was particularly apparent among transit amplifying cells and enterocyte progenitors **(figure 2H, S3B)**. Flow cytometry validated that stretch increased the population of Ki67-positive cells lacking expression of the stem cell marker Lgr5, which presumably represent proliferative progenitor cells **(figure S3C)**. As stretch application did not consistently increase the number of progenitor cells across both replicates **(figure 2E)**, mechanical strain may particularly alter their cell cycle entry and/or exit without affecting their abundance. Finally, whereas most cell types showed a substantial number of differentially expressed genes following stretch, the largest number of transcriptional changes were identified in enterocytes and their progenitors **(figure S4A-B**). As these transcriptional responses were particularly pronounced at 15% stretch **(figure S4A)**, we focused in further analyses on this condition. Altogether, our unbiased transcriptomic analyses uncovered that mechanical strain triggers a decrease in enterocytes, while enhancing the proliferation rate of precursor populations.

### Mechanical strain shifts absorptive lineage composition toward an early enterocyte state

The observed changes in enterocyte abundance were not attributable to increased cell death (**figure S2A)**. Given that cell loss from the intestinal epithelium can result from both apoptotic and live cell extrusion (Eisenhoffer et al., 2012; Krueger et al., 2025), we further performed live imaging of organoids under cyclic mechanical stretch, which revealed no effect on the rate of cell extrusion (**figure S2B-C**). Closer examination of the abundance of enterocytes revealed that this cell population was heterogeneously affected by mechanical strain **(figure 2C)**. As was particularly apparent at 48 hours, mechanical strain exclusively reduced a specific part of the enterocyte population, while another subset became more abundant **(figure 2C, S5A)**. To further characterize these differential effects of strain across the enterocytes, we performed unsupervised clustering and compared the abundances of each enterocyte subpopulation in response to 15% stretch **(figure 3A, S5B)**. Indeed, one of the enterocyte subpopulations (EC cluster 0) consistently decreased in organoids subjected to strain, while a transcriptionally distinct enterocyte population (EC cluster 1) increased in abundance **(figure 3B, S5C)**. The effects on enterocyte subcluster 2 were inconsistent across biological replicates, and this cluster was therefore not further examined **(figure 3B, S5C)**. These findings indicate that mechanical strain triggers a shift between enterocyte subpopulations with distinct transcriptional profiles, rather than uniformly reducing enterocyte numbers.

**Figure 3.**
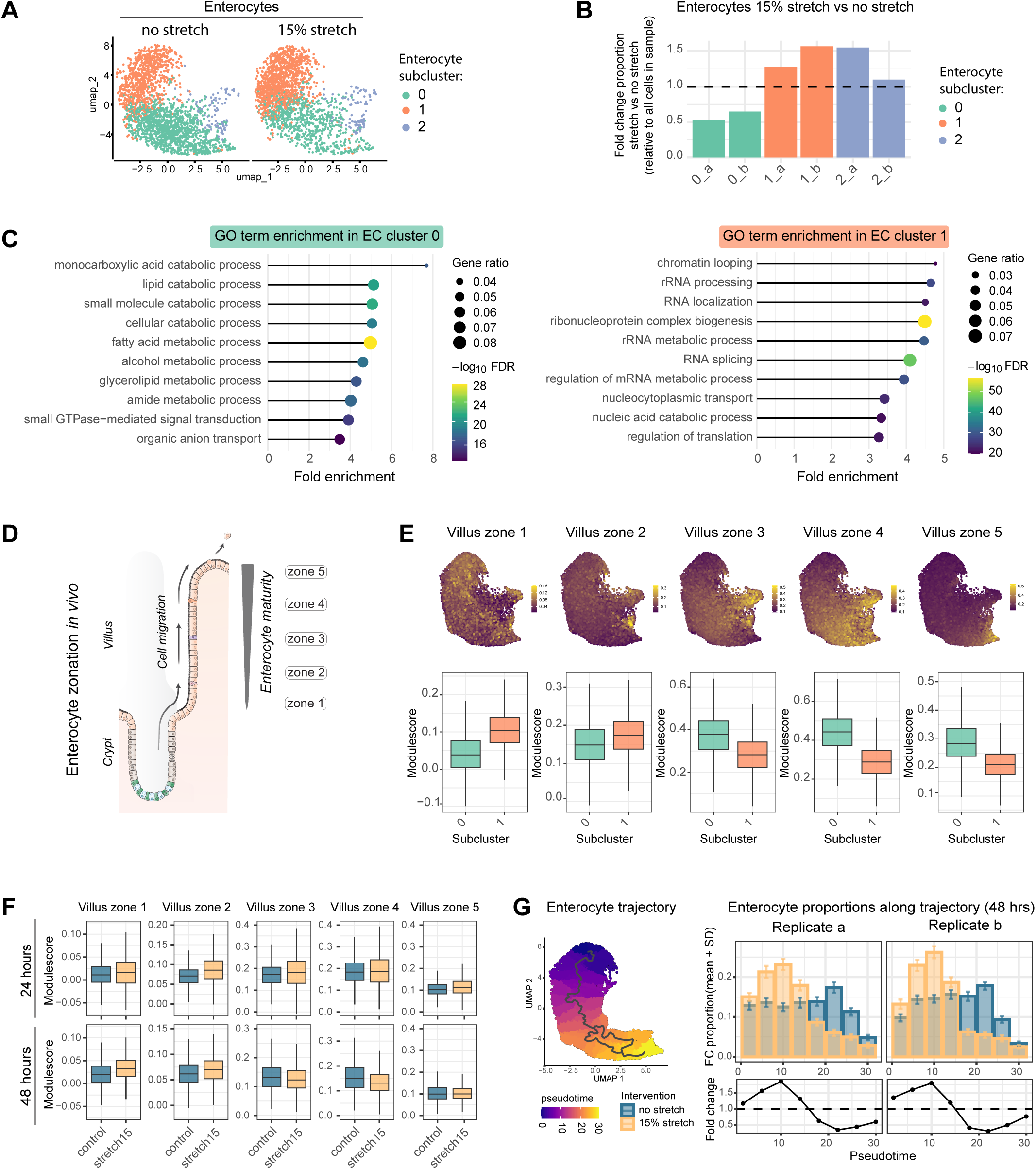
Mechanical strain triggers a shift from late to early-stage enterocytes. **(A)** UMAP of enterocytes from organoids with or without 48 hours of 15% cyclic stretch. Unsupervised clustering identified three enterocyte subclusters present under both conditions. Data includes both biological replicates. (**B**) Proportion of enterocyte subclusters after 48 hours of 15% stretch, relative to their respective non-stretched controls. Bars of the same color represent individual replicates**. (C)** GO term enrichment analysis (Biological Process) based on marker genes from enterocyte subclusters 0 and 1. Analysis is performed on pooled stretched and non-stretched data. (**D)** Schematic representation of in vivo intestinal enterocyte zonation along the villus axis, based on Moor et al. (Moor et al., 2018). (E) Top: UMAPs of enterocytes (15% stretched and non-stretched at 48 hours, pooled from both replicates) colored by module scores for each villus zonation gene signature (Moor et al., 2018). Bottom: boxplots comparing zonation module scores between enterocyte subclusters 0 and 1. (**F**) Boxplots comparing zonation module scores across all enterocytes with and without 15% stretch. Data represent values from both biological replicates. **(G**) Left: UMAP showing pseudotime progression across enterocytes inferred by trajectory analysis (Monocle 3), highlighting a branch from subcluster 1 to 0 (pooled data from all stretch conditions, replicates, and timepoints). Right: Distribution of enterocytes along pseudotime bins for non-stretched and 15% stretched conditions at 48 hours. Mean and SD were calculated using bootstrapping.

To identify what distinguishes the enterocyte subpopulations that respectively decreased (cluster 0) and increased (cluster 1) in response to strain, we performed GO term enrichment analysis on genes that were enriched in either population **(figure 3C, S5D)**. In enterocyte cluster 0, the top enriched processes were all related to nutrient absorption and metabolic activity **(figure 3C)**. In contrast, enterocyte cluster 1 showed enrichment of processes associated with transcription and translation **(figure 3C)**. The transcriptional landscapes of both strain-responsive enterocyte populations are thus associated with distinct cellular functions, and their strain-induced shift in abundances may therefore adjust specific absorptive lineage functions.

Previous single-cell transcriptome analysis of the mouse intestinal epithelium uncovered that enterocytes adopt functionally distinct transcriptional states during their migration along the villus axis **(figure 3D)** (Moor et al., 2018). Specifically, enterocytes at the villus base (zone 1-2) are enriched in programs driving gene transcription, translation and mitochondrial activity. In contrast, more mature enterocytes located higher up in the villus gradually acquire nutrient-absorbing functions, including uptake of carbohydrates (zone 3), peptides (zone 4), and lipoprotein biosynthesis (zone 5). Interestingly, these functionally distinct enterocyte states resemble the transcriptional profiles that distinguished the two stretch-responsive enterocyte populations in our organoids **(figure 3C)**. Indeed, projecting these villus zonation signatures onto our organoid data revealed that upper villus markers (zone 3-5) were enriched in the organoid enterocyte population that decreased with stretch (cluster 0), while lower villus signatures (zone 1-2) were enriched in the stretch-expanded enterocyte population (cluster 1) **(figure 3E)**. Moreover, independently of enterocyte sub-clustering, lower villus signatures were enriched in enterocytes derived from stretched organoids, whereas non-stretched enterocytes showed enrichment for mid-and upper-villus signatures (zone 3-4) **(figure 3F).** Notably, the enrichment of lower villus signatures in stretched enterocytes was already evident at 24 hours, whereas the reduction of upper villus signatures only became apparent at 48 hours **(figure 3F)**, suggesting that the increase of enterocytes representing lower villus-like cells precedes the loss of upper villus-like enterocytes. Finally, trajectory analysis of the enterocytes **(figure S5E-F)** confirmed that strain increased the number of early-stage enterocytes at the expense of late-pseudotime enterocytes **(figure 3G)**. This indicates that mechanical strain redistributes the enterocyte lineage towards a less mature state that transcriptionally resembles bottom-villus enterocytes.

### Mechanical strain expands early enterocytes at the crypt-villus interface

To validate the strain-induced shift from late to early enterocytes beyond our single cell transcriptome analyses, we utilized individual marker genes of the different enterocyte villus zones **(figure 4A)**. All selected zonation markers showed the expected expression pattern across the early-and late-stage enterocyte populations in our organoids, and exhibited corresponding strain-induced expression changes in our scRNA-seq analysis **(figure S6A-B)**. Note that marker genes for zone 1 were not restricted to the enterocyte population but were broadly present in stem and progenitor cells as well, and were therefore not included for further analyses **(figure S6C)**. Next, subjecting organoids to 15% cyclic stretch and subsequent RT-qPCR analysis for the selected zonation marker genes confirmed the significant downregulation of genes marking higher villus enterocytes in stretched organoids (zone 3: *Fabp1*, *Adh1*, *Gda*. Zone 4: *Treh*, *Xdh*, *Ephx*2. Zone 5: *Apoc3*, *Apoc4*) **(figure 4B)**. In contrast, the zone 2 marker *Ak2* was significantly upregulated by strain **(figure 4B)**.

**Figure 4.**
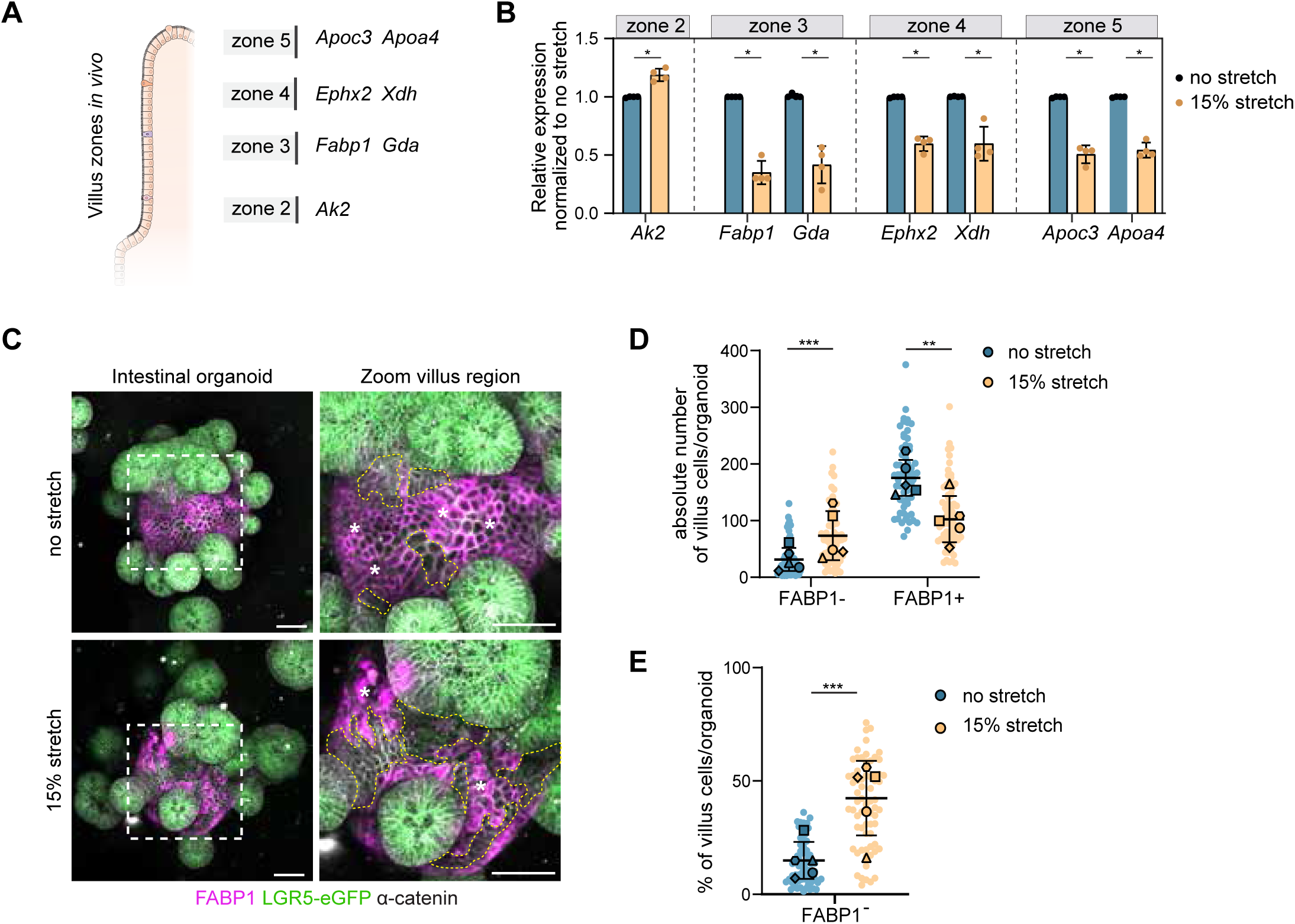
Mechanical strain promotes expansion of FABP1neg early enterocytes at the crypt-villus interface. **(A)** Schematic representation of genes marking villus zones, identified in Moor et al. (Moor et al., 2018). **(B)** RT-qPCR of selected zonation marker genes in small intestinal organoids with or without 15% cyclic stretch for 24 hours. Data are shown as mean ± SD from 4 independent experiments, relative to non-stretched controls. *p < 0.05; unpaired Mann-Whitney test on ΔΔCT values per gene. **(C)** Representative maximum intensity z-projection images of organoids cultured with or without 15% cyclic stretch for 24 hours, immunostained for FABP1. FABP1-positive cells mark mature enterocytes (villus zone 3; asterisk), whereas FABP1-negative villus cells represent early-stage enterocytes (outlined by dashed line). Scale bar: 50 μm. **(D)** Quantification of FABP1-high and FABP1-low cells per villus region in individual organoids with or without 15% stretch. Each point represents one organoid (n = 64 without stretch, n = 67 with stretch, from 5 independent experiments), with larger points representing replicate means, and horizontal lines the overall mean ± SD. Statistics: Negative binomial generalized linear mixed-effects models modeling the absolute count of FABP1-low and-high cells with random intercepts and slopes for condition across replicates. ** p < 0.01, *** p < 0.001 **(E)** Percentage of FABP1-high and FABP1-low cells per villus region in individual organoids from figure 4D. Each point represents one organoid (n = 64 without stretch, n = 67 with stretch, from 5 independent experiments) with larger points representing replicate means, and horizontal lines the overall mean ± SD. Statistics: Binomial generalized linear mixed-effects

To corroborate whether these transcriptional changes correspond to the redistribution of late-to-early-stage enterocytes, we visualized the distribution of both populations across the organoids by immunostaining for specific enterocyte marker proteins. First, we co-stained the late enterocyte marker FABP1 (zone 3) together with the general enterocyte marker KRT20, which was expressed throughout enterocytes and only mildly decreased in response to strain **(figures S6D-E)**. This established that the majority of KRT20-positive enterocytes in organoids expressed high levels of FABP1 protein, consistent with a mature enterocyte state **(figure S6F)**. A subset of villus cells located near the crypt boundary, which already began to show KRT20 expression, lacked detectable FABP1 protein **(figure S6F)**. This cell population also lacked KI67 expression **(figure S6G)** and thus represents early-stage enterocytes that have exited the crypt, but did not yet transition to a fully mature state. Following 24 hours of 15% cyclic stretch, this population of FABP1-negative villus cells had markedly expanded **(figure 4C)**, accompanied by a reduction in absolute FABP1-positive villus cells per organoid **(figure 4D-E)**. This shift occurred across crypt-villus border regions with different orientations, suggesting that this does not require a specific stretch direction. Collectively, our transcriptional and immunostaining analysis indicate that mechanical strain alters enterocyte lineage composition by halting maturation of early-stage enterocytes at the villus base.

### Strain controls upstream regulators of enterocyte maturation across intestinal cell populations

Since the generation of mature enterocytes is driven by induction of specific transcriptional programs (Kolev & Kaestner, 2023; Noah et al., 2011), we next investigated how mechanical strain influences the activity of transcriptional regulators. By mapping genes that were significantly up-or downregulated in stretched organoids to their known upstream transcriptional regulators, we inferred regulators with altered activity in response to mechanical strain **(figure 5A, S4A)**. Assessing upstream regulator activity across each cell type of the absorptive lineage revealed several transcriptional (co-)factors with stretch-induced increased or decreased activity **(figure 5B, S7A)**. This included activation of the well-established mechanoresponsive transcriptional co-activator YAP across all cell types in stretched organoids **(figure 5B, S7A)**. Other transcriptional regulators of which target genes were globally upregulated across stretched cell types included Foxa2, Rela, Tcf7L2, Klf4, and Nfat5 **(figure 5B, S7A)**. Among the upstream transcriptional regulators most prominently inhibited by strain, we identified multiple major drivers of enterocyte differentiation *in vivo*, including Hnf4α (Babeu et al., 2009; Cattin et al., 2009; Chen, Toke, Luo, Vasoya, Fullem, et al., 2019), Hnf4γ (Baraille et al., 2015; Chen, Toke, Luo, Vasoya, Aita, et al., 2019; Lindeboom et al., 2018), and Cdx2 (Benoit et al., 2010; Gao et al., 2009; Hryniuk et al., 2012; San Roman et al., 2015) **(figure 5B, S7A)**. Importantly, the reduced activity of these transcriptional regulators was not only apparent in enterocytes, but extended to immature precursor cell types (stem and TA cells) **(figure 5B-5C)**. Comparing Hnf4α activity across these populations at different stretch durations revealed a temporal sequence: Hnf4α activity was first suppressed by strain within absorptive progenitor populations (stem cells, TA and enterocyte progenitors) at 24 hours, and only later led to a prominent reduction of Hnf4α target gene expression in differentiated enterocytes at 48 hours **(figure 5B-C)**. Mechanical strain thus interferes with the main transcriptional programs driving enterocyte differentiation early on in the absorptive lineage. Together, our findings support a model in which mechanical strain attenuates key transcriptional programs involved in enterocyte maturation, thereby arresting absorptive cells in an immature state and ultimately limiting the production of fully differentiated, nutrient-absorbing enterocytes.

**Figure 5.**
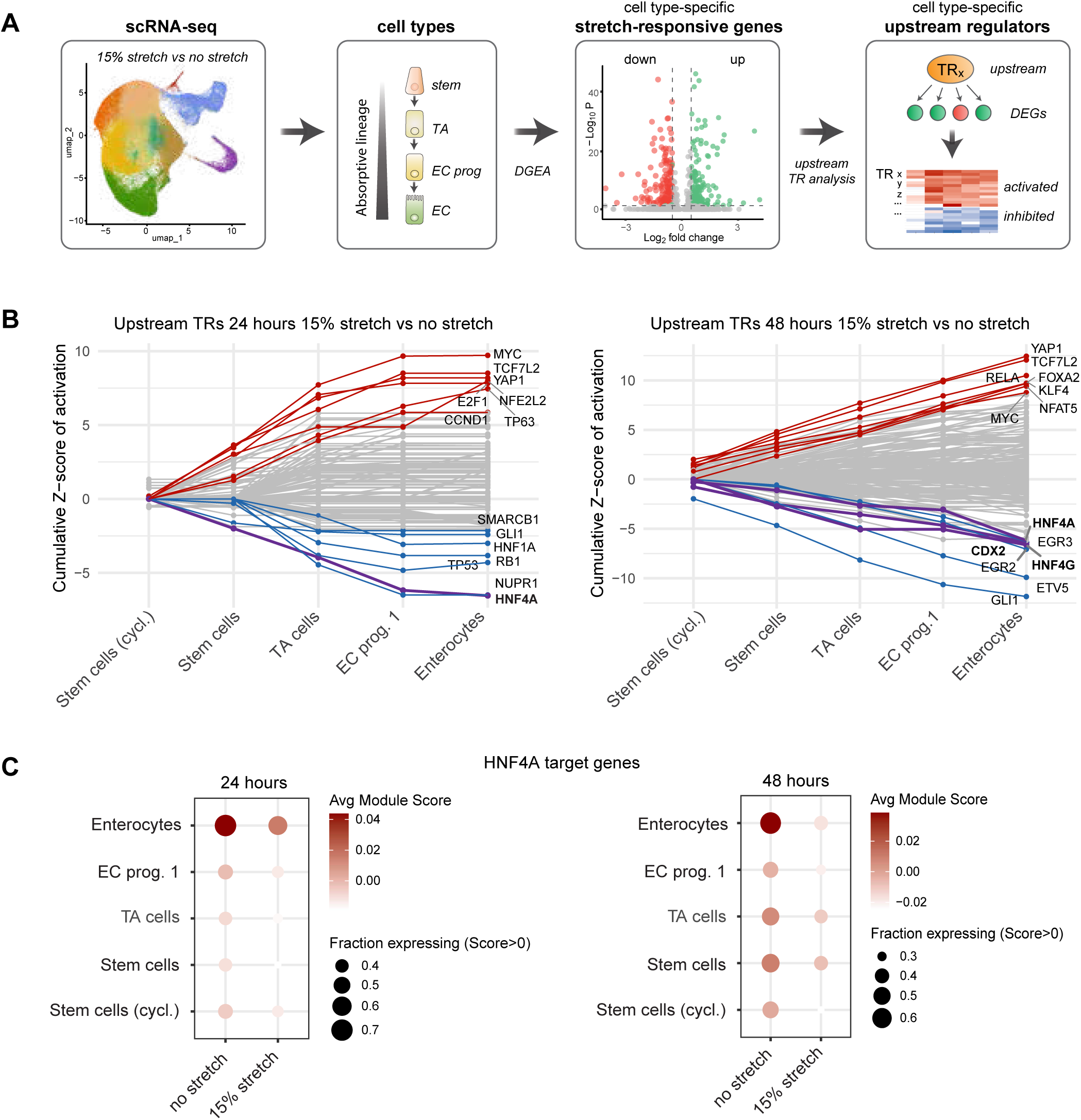
Transcriptional regulator identification upstream of strain-responsive transcriptomes per cell type. **(A)** Schematic representation of workflow to identify upstream transcriptional regulators of stretch-responsive differentially expressed genes. Differentially expressed genes between 15% stretch and no stretch for each cell type of the absorptive lineage are used to predict the activation level of upstream transcription regulators whose, based on known regulator-target gene interactions through literature. **(B)** Line plots showing cumulative activation z-scores of upstream regulators across the cell types of the intestinal absorptive lineage after 15% stretch for 24 hours (left) and 48 hours (right). Top 7 activated and inhibited regulators are highlighted. Average Z-score is calculated from both biological replicates. **(C)** Dot plot showing module scores of Hnf4α target gene expression, based on Gunewardena et al., 2022, across the cell types of the intestinal absorptive lineage with and without 15% stretch for 24 and 48 hours.

## Discussion

In this study, we investigated the influence of mechanical forces on the cellular composition of the intestinal epithelium by external stretch application to intestinal organoids, combined with high-throughput single-cell transcriptomics. We found that mechanical strain shifts the absorptive cell lineage towards a less mature state by expanding early-stage enterocytes and concomitantly reducing transcriptionally distinct, late-stage enterocytes. Our findings imply that the intestinal epithelium adapts its absorptive functions in response to mechanical input by modulating the maturation status of the absorptive lineage, revealing a novel concept of force-sensitive regulation of intestinal cell type composition.

Our data support a model in which mechanical strain halts the maturation of early-stage enterocytes and thereby limits their transition towards specialized nutrient-absorbing cells. This is consistent with the observed increase in abundance of early enterocytes at the base of the villus **(figure 4D-E)**, as well as suppression of transcriptional regulators that promote enterocyte maturation, including Hnf4α, in enterocyte precursor populations **(figure 5C, S7A)**. The observed expansion of early enterocytes occurs in absence of increased apoptosis or cell extrusion in stretched organoids (**figure S2A-C**), which argues against enterocyte loss driving the observed shift in enterocyte populations. We cannot fully exclude that reversion of late enterocytes into a less mature state contributes to the strain-induced shift between enterocyte populations. However, although differentiated intestinal cells have been shown to dedifferentiate to a stem-like state during intestinal regeneration (Hageman et al., 2020), specific dedifferentiation of late-stage enterocytes into immature enterocytes has not yet been observed.

Identification of mechanoresponses in organoid models may ultimately provide insights into the complex roles of mechanical forces in the native intestine. One model emerging from our findings is that enterocyte zonation along the villus axis can be influenced by intrinsic tensile forces along the epithelium. Cell migration towards the villus tip generates a gradient of tension that peaks at the villus base, and thus anti-correlates with enterocyte maturity along the villus (Krndija et al., 2019; Moor et al., 2018). Potentially, tension levels within each villus zone may locally instruct enterocyte fate, with the tension gradient cooperating with signaling gradients such as BMP in regulating zonation (Beumer et al., 2022). Ectopically stretching organoids may sustain elevated tension levels, thereby preventing enterocytes from progressing to a mature state. These insights warrant further experimental investigation into to contribution of forces to intestinal villus zonation, which would require *in vivo* manipulation of forces experienced by the intestinal epithelium. Such studies may reveal how force-dependent modulation of enterocyte maturation influences the absorptive functions of the intestine. Moreover, they may uncover to what extent forces could potentiate the adaptation of zonation patterns across different regions of the intestine (Zwick et al., 2024).

Our study identified strain-induced changes in the most abundant lineage of the intestinal epithelium. However, our data do not exclude that more rare intestinal cell types may similarly be modulated by forces, which may be more difficult to capture with our current analyses. Mechanical forces and downstream mechanosensing have previously been implicated in diverse other cellular processes in the intestine, including stem cell proliferation and maintenance (Baghdadi et al., 2024; Y. Li et al., 2020; Meng et al., 2022), secretory versus absorptive fate decisions (Baghdadi et al., 2024), cell extrusion (Krueger et al., 2025), and secretory cell functions (Alcaino et al., 2018; Fang et al., 2023; Treichel et al., 2021; Xu et al., 2021). Most of these previously described intestinal mechanoresponses have been linked to the mechanosensitive calcium channels Piezo (Alcaino et al., 2018; Baghdadi et al., 2024; Fang et al., 2023; He et al., 2018; Tallapragada et al., 2021; Treichel et al., 2021; Xu et al., 2021). Elucidating the mechanosensor that couples mechanical strain to enterocyte maturation, and defining how this is functionally coordinated with other mechanoresponses across intestinal cell types to orchestrate tissue-level regulation, represents an important direction for future studies

## Materials and Methods

### Antibodies and reagents

The following antibodies were used at the indicated concentrations for immunofluorescence (IF) and flow cytometry (FC): Fabp1 (D2A3X XP Cell Signaling, 13368S, IF 1:200), α-catenin (15D9, Enzo, ALX-804-101-C100, IF 1:200), Cytokeratin-20 (Dako, M7019, IF 1:50), Cleaved Caspase-3 D175 (Cell Signaling, CS9661, IF 1:200), Ki67 (Abcam, 15580, FC 1:200) and Alexa Fluor conjugate secondary antibodies (Life Technologies, IF 1:500, FC 1:500). The following dyes were used at the indicated concentrations: DAPI (Sigma, SKU: D9564, IF 1:1000) and DRAQ7 (Cell Signaling, 7406S, FC 1:200).

### Organoid culture

Organoids isolated from the jejunum and ileum of Lgr5-DTR-eGFP mice (Tian et al., 2011) were a kind gift from Katharina Sonnen (Hubrecht institute, The Netherlands). Organoids were embedded in basement membrane extract (Cultrex RGF BME Type 2, Bio-techne, 3533-005-02) and cultured in ENR medium containing Advanced DMEM/F12 (Gibco, 12634-010), supplemented with 10 mM HEPES (Gibco, 15630-056), 1x Glutamax (Gibco, 35050-038), 1% Penicillin-Streptomycin (Merck, P0781-100ML), 10% Noggin-conditioned medium, 5% R-spondin1-conditioned medium, 1x B27 supplement (Thermo Fisher, 17501-001), 1 mM N-Acetyl Cysteine (Sigma, A9165-5G) and 50 ng/ml human recombined epidermal growth factor (EGF, Peprotech, AF-100-15). Organoids were passaged every 4-7 days by removing them from BME, mechanical dissociation into crypt fragments, and embedding in fresh BME. Medium was refreshed every 2-3 days. Culture conditions were kept stable in the incubator at 37°C with 5% CO_2_. Organoids with stable expression of H2B-mScarlet-I were generated by lentiviral with virus containing the pLV hEF1α-H2B-Scarlet-I-IRES-blast plasmid (generated by In-Fusion cloning) and selection based on Blasticidin resistance.

### Stretch application to intestinal organoids

Poly-dimethyl siloxane sheets (PDMS-E, 600 µm thickness, SolSep B.V., HFS-PDMS-06) were functionalized through plasma treatment (Harrick Plasma, PDC-001) followed by 1 hour incubation with 10% APTES in ethanol at 60°C. Sheets were then washing three times with 100% ethanol and two times with PBS for 10 minutes each, completed drying for 30 minutes at 60°C and 25 minutes incubation with 1.5% glutaraldehyde in PBS at room temperature. Sheets were washed 5 times with PBS for 10 minutes on a shaker to get rid of unbound glutaraldehyde and dried for 30 minutes at 60°C. To seed organoids, functionalized sheets were mounted to the MechanoCulture TM uniaxial stretch device (CellScale Biomaterials Testing) after UV-and ethanol-based sterilization of the device.

Organoids were seeded as for general culturing. Briefly, 4-6 days old, differentiated organoids were dissociated into crypt segments, recovered by centrifugation at 250 x g, and embedded in 20 μl BME domes that were plated on the PDMS surface. Incubation at 37°C for 30 minutes enabled BME polymerization and covalent attachment of BME to the PDMS. Organoids were grown in ENR medium at 37°C and 5% CO_2_, and medium was refreshed after 3 days. Application of mechanical stretch was started 4 days after seeding after organoids had budded into 3D compartmentalized structures containing all specialized cell types. The indicated levels of uniaxial stretch were applied to the PDMS sheet at a frequency of 0.1 Hz (10 seconds period, ramp program) through motor-based actuator movements controlled by CellScale software.

### Measuring whole-organoid strain

To determine whole-organoid strain **(figures 1B-E, S1B-C, S1E)**, images of Lgr5-DTR-eGFP mouse intestinal organoids expressing H2B-mScarlet-I were acquired before and during application of static stretch by mounting the CellScale stretch device on a Zeiss Cell Observer widefield microscope equipped with Orca Flash 4.0 camera (Hamamatsu), using a 10x (NA = 0.3) or 20x (NA = 0.75) objective and NIS Elements software. In all other experiments, organoid strain was confirmed based on brightfield images before and during static stretching acquired on an EVOS-M5000 widefield microscope, using a 2x (NA = 0.08) or 4x (NA = 0.16) objective. Images were processed using ImageJ software, and whole-organoid strain percentages were calculated based on change in absolute length of the entire organoid along the stretch axis (axial strain, Ɛ_xx_) or parallel to the stretch axis (perpendicular strain, Ɛ_yy_).

### Live imaging during stretch application and local strain measurements within organoids

To identify local intercellular strain patterns within organoid compartments, Lgr5-DTR-eGFP mouse intestinal organoids expressing H2B-mScarlet-I were imaged live before and during stretch on a Nikon Spinning Disk Confocal Microscope (Yokogawa CSU-W1) using a 20x dry objective (NA = 0.75) and NIS Elements software. Z-stack images of H2B-mscarlet-I, Lgr5-DTR-eGFP and brightfield were acquired with 2.5 μm step size.

To overlay stretched and non-stretched images, manual image registration (Brown, 1992) and analysis was performed using a custom-written plugin for OrganoidTracker (Kok et al., 2020). Cells with defined features with and without stretch were used as manually annotated 3D landmark points, and used to construct a registration map (with resolution of 1 x 1 x 1 μm) that maps every voxel from the pre stretch to the stretched z-stack image. The registration map was constructed from the landmarks using linear interpolation using the function scipy.interpolate.griddata from Scipy (Jones et al., 2001). The registration map was smoothed using a Gaussian filter (scipy.ndimage.gaussian_filter) with *σ* = 15 μm.

To visualize axial strain levels within organoid regions **(figure 1F)**, we compared the point 5 μm lower on that axis to the point 5 μm higher, and measured how far those points were apart on that axis during stretch according to the registration map. In absence of strain, the points will be 10 μm apart. Strain was calculated as:

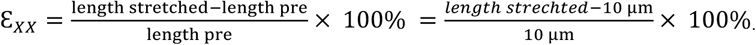

Color-coded axial strain levels were projected on the maximum z intensity projection of the organoid.

To determine axial strain levels of individual crypt and villus compartments **(figure 1G)**, the distance between two points at the edges of individual crypt or villus regions was manually measured parallel to the stretch axis, before and during application of 30% static stretch:

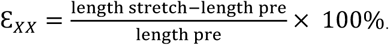

To determine live-cell extrusion rates in intestinal organoids **(figure S2B-C)**, Lgr5-DTR-eGFP mouse intestinal organoids expressing H2B-mScarlet-I were imaged live in the absence or presence of 15% cyclic stretching (0.1 Hz) in a temperature controlled chamber at 37°C and 5% CO_2_. Imaging was performed on a Nikon spinning disk confocal microscope (Yokogawa CSU-W1) using a 20x dry objective (NA = 0.75) and NIS Elements software. A 70 μm z-stack with 2.5 μm step size was acquired every 10 minutes when organoids were in their non-stretched (rest) position. Image stacks were registered and stabilized using the Fast4Dreg plugin in ImageJ. Extrusion events were quantified manually from color-coded maximum-intensity z-projections generated in ImageJ.

### Single-cell RNA sequencing

Organoids subjected to mechanical strain for 24 or 48 hours were dissociated from the BME by fragmentation and subsequent incubation with ice-cold Cell Recovery Solution (Corning, 354253) for 20 minutes. Organoids were washed twice in advanced DMEM/F12, recovered by cold centrifugation at 250 x g for 5 minutes, and snap frozen as whole organoid pellets in liquid Nitrogen, followed by storage at-70°C. Next, organoids were fixed in 4% Formaldehyde in Fix & Perm Buffer (10x Genomics PN-2000517) for 24 hours at 4 °C without agitation, then washed with chilled PBS and quenched in Quenching Buffer (10x Genomics PN-2000516). For dissociation, organoids were incubated in 0.2 mg/ml Liberase at 37 °C for 20 minutes, followed by manual trituration. Cell suspensions were filtered (30 μm), washed, resuspended in Quenching Buffer, and counted using an automated cell counter (Countess II/Cellaca MX). Samples were then processed for multiplexed single cell RNA-sequencing (10x Genomics) on a NovaSeq X Plus Illumina system by Single Cell Discoveries following the manufacturer’s protocol.

### Single-cell RNA sequencing data analyses

Raw sequencing data (fastq files) were processed and mapped with CellRanger software (version 7.1.0) using Cellranger multi pipeline. 10X mouse genome reference (Mus Musculus; refdata-gex-mm10-2020-A; GRCm38, Ensembl 98) and respective 10X probe set reference csv file (10X Flex probe set Chromium_Mouse_Transcriptome_Probe_Set_v1.0.1_mm10-2020-A.csv) were provided to Cellranger multi pipeline for read alignment. Subsequent processing and analysis were performed in R (version 4.4.3) using Seurat (version 5.2.1). Briefly, cells were selected based on the presence of at least 500 transcripts and 300 genes, a log10 (genes-per-transcript) value greater than 0.8, and a mitochondrial gene read fraction below 0.1. Doublets were identified using scds (version 1.22.0) with a hybrid doublet score threshold of 1.8. Following cell filtering, data were normalized using sctransform v2 in Seurat and clustered with the Leiden algorithm.

Differentially expressed genes for each cluster were identified using a Wilcoxon rank-sum test, considering genes with a minimum log2(fold change) of 0.25. To compare organoid cell clusters to primary cell types, we identified common marker genes (P < 0.01 and log2(fold change) > 1.01) between organoid clusters and primary mouse intestinal cell clusters as defined in a published scRNA-seq dataset (Haber et al., 2017). The significance of the overlap in marker genes was assessed using a binomial test, with probabilities estimated by randomizing the primary tissue marker gene list across 10,000 iterations. Next, differential abundance testing across defined clusters was conducted using propeller (speckle version 1.6.0), while local neighborhood differential abundance was assessed using MiloR (version 2.2.0).

Analyses of enterocyte subtypes were performed by subsetting and reprocessing the enterocyte population following the same pipeline. GO-term enrichment analysis between enterocyte subclusters was carried out using clusterProfiler (version 4.16.0). Trajectory inference in enterocytes was performed using Monocle3 (version 1.3.7) and mean ± SD cell proportions across pseudotime were calculated by bootstrapping 100 replicates.

For differential gene expression analysis between conditions, each replicate was analyzed separately using MAST (version 1.32.0). Genes with significantly altered expression (adjusted P < 0.05) were subsequently used to identify upstream regulators based on literature and existing databases using Ingenuity Pathway Analysis software (Qiagen). Upstream regulators were filtered for significant transcriptional regulators with a z-score of activation in both biological replicates.

### RT-qPCR

Organoids were isolated from BME by fragmentation in cold advanced DMEM/F12 medium, recovered by cold centrifugation at 250 x g for 5 minutes, and subsequently lysed in RLT lysis buffer (Qiagen, 79216). Total RNA was isolated using the RNeasy kit (Qiagen) with DNase treatment (Qiagen) according to the manufacturer’s instructions, and cDNA was synthesized using the iScript^TM^ cDNA synthesis kit (Bio-Rad, 1708891). RT-qPCR was performed using PowerTrack SYBR Green Master Mix (Fisher Scientific, 16555231) and run on QuantStudio5 (Thermo Fisher Scientific). Technical triplicates were included for each sample. Fold changes were determined using the 2-ΔΔCT method, with normalization to pooled expression of housekeeping genes (*Pbgd, Hnrnpa, Cyca, Tuba1*). Optimized forward and reverse primers used for each gene are listed below.

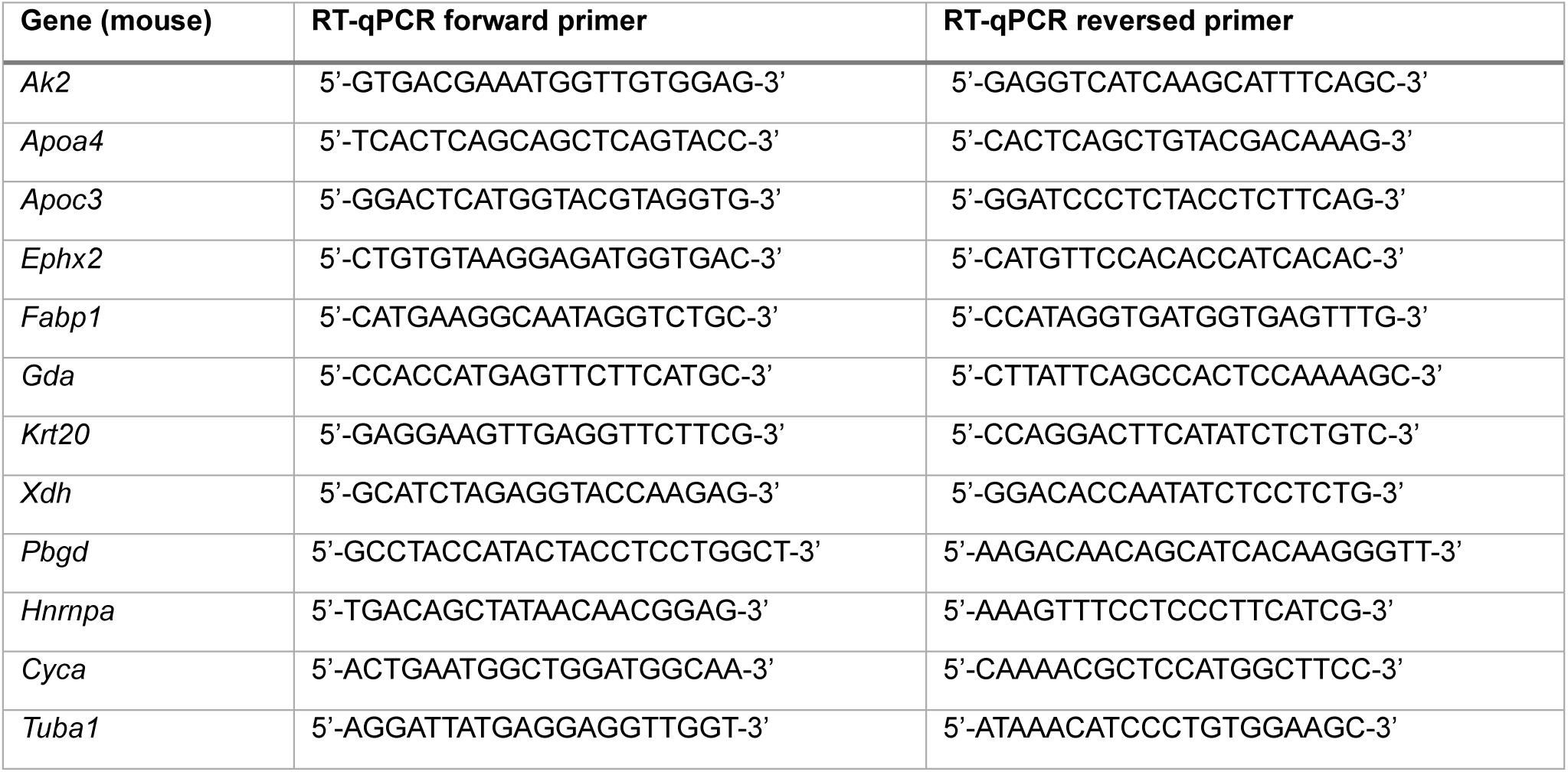

### Immunofluorescence

Intact BME droplets with stretched or non-stretched organoids were transferred to microcentrifuge tubes (Eppendorf) and fixed with 4% PFA and 0.25% glutaraldehyde in PBS for 20 minutes at room temperature in the dark. Glutaraldehyde-derived reactive aldehyde groups were quenched by washing three times 10 minutes with 1% Sodium Borohydride in PBS, and washing three times 10 minutes with PBS. Organoids were permeabilized and blocked with 1% BSA, 10% DMSO and 2% Triton-X100 in PBS (PBD2T) for at least 4 hours at 4°C, and incubated with primary antibodies in PBD2T at 4°C overnight. After washing with PBS, secondary antibodies in PBD2T were incubated overnight. Z-stack images of 90 μm with 5 μm step size were acquired at a Zeiss LSM880 laser-scanning confocal microscope using a 20x objective (NA 0.75). Images are presented as MAX intensity z-projections and were processed using ImageJ software.

### Flow-cytometry

Organoids were isolated from BME by fragmentation in cold advanced DMEM/F12 medium, recovered by cold centrifugation at 250 x g for 5 minutes and dissociated into single cells by TrypLE treatment (Life Technologies, 12605028) at 37°C, supplemented with 10 µM Y-27632 dihydrochloride (Gentaur, 607-A3008-50MG). After washing with PBS containing 1 mM EDTA and 1% FCS, cells were fixed and permeabilized overnight at 4°C in 70% ethanol. Cells were blocked and incubated with primary and secondary antibodies in PBS containing 1% BSA and 0.05% Tween 20 (Merck, 8221840050) for 1 hour in the dark in at room temperature, and washed with PBS with 1% BSA. Flow cytometry was performed using the BD FACS Celesta machine, and data were analyzed using Cytobank.

## Statistical analyses

Single cell RNA sequencing analyses and statistics were performed in RStudio. Statistical analyses of organoid strain levels, immunofluorescence, RT-qPCR and flow-cytometry were performed in GraphPad Prism (version 10).

## Data availability

Single cell RNA sequencing data will be made publicly available at the Gene Expression Omnibus (GEO) upon publication.

## Acknowledgements

We thank the members of our laboratories for helpful discussions and Willem-Jan Pannekoek and Danijela Matic Vignjevic for critical reading of the manuscript. We thank Jesse Bosma (UMC Utrecht) for designing and 3D printing control stretch devices. This work was supported by the Netherlands Organisation for Scientific Research (NWO; 016.Vidi.189.166), the Dutch Cancer Foundation (KWF-15290A), the EU/EFPIA Innovative Medicines Initiative Joint Undertaking (PERSIST-SEQ, grant No 101007937) and NWO Gravitation Project: BRAINSCAPES: A Roadmap from Neurogenetics to Neurobiology (NWO: 024.004.012).

## Author contributions

R.M.H., B.v.S., A.v.O., and M.G. conceived the study. R.M.H. and B.v.S. designed the experimental approach. R.M.H. performed the organoid experiments. B.v.S. conducted the scRNA-seq data analysis. K.v.d.A. helped with organoid experiments. M.J.V. performed qPCR experiments. R.M.H., B.v.S., K.v.d.A., M.J.V., and M.C.v.d.N. analyzed the data. R.K. and M.R.C. performed and supervised the imaging analysis of intra-organoid strain. R.M.H. and B.v.S. assembled the figures. R.M.H. and M.G. drafted the manuscript, B.v.S. and A.v.O. edited the manuscript. M.G. and A.v.O. supervised the research and acquired funding.

## Declaration of generative AI and AI-assisted technologies in the manuscript preparation process

During the preparation of this work, the authors used ChatGPT (openAI) to improve grammar and readability. After using this tool, the authors reviewed and edited the content as needed and take full responsibility for the content of the publication.

---

**Figure S1.**
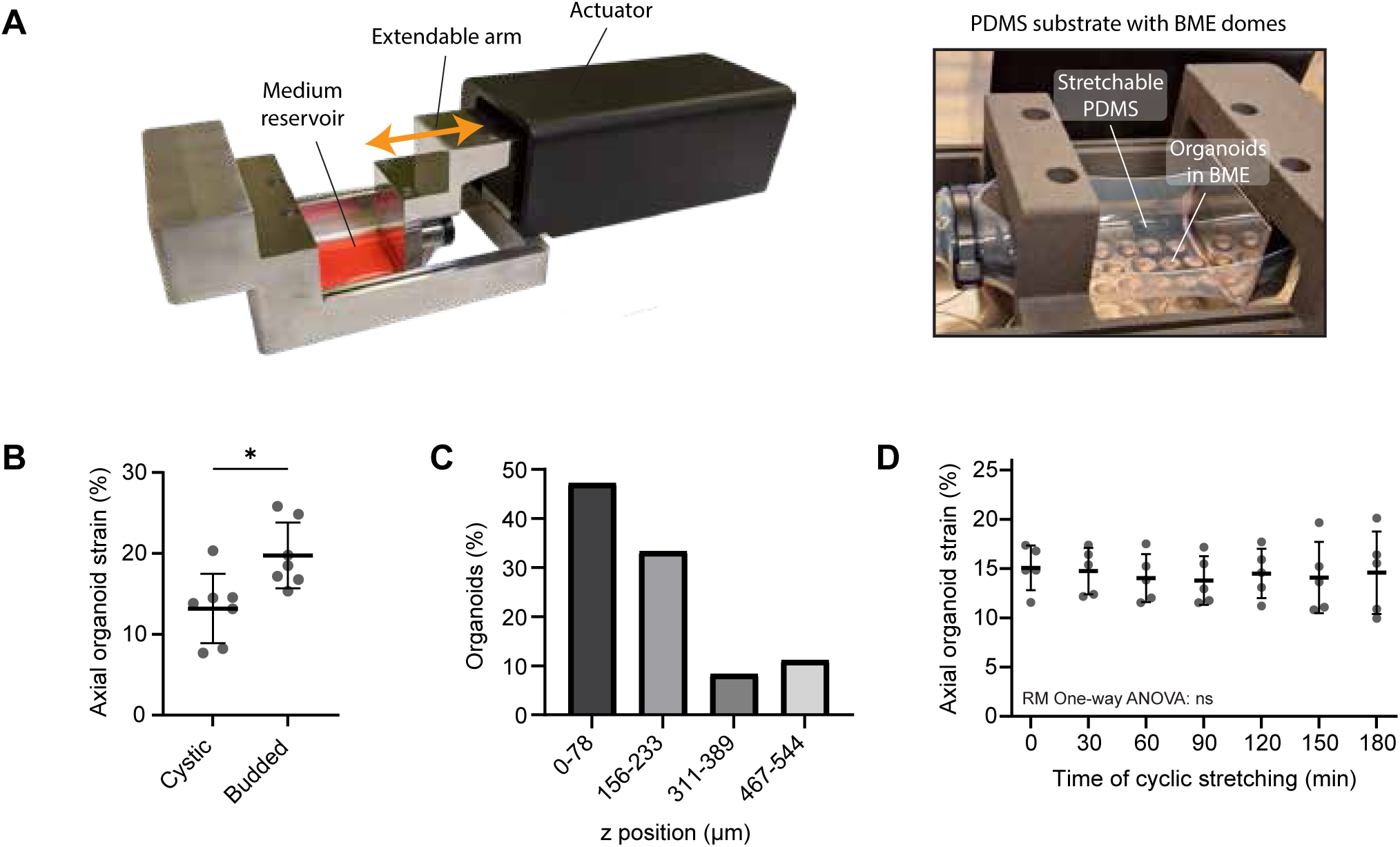
Mechanical stretch application to intestinal organoids. **(A)** Stretch device used to apply uniaxial, cyclic mechanical strain to intestinal organoids embedded in BME domes coupled to underlying PDMS. **(B)** Axial strain (%) of cystic versus budded (crypt-containing) organoids during 30% static stretch. Analyzed organoids were located at comparable z-heights from the PDMS surface (within 155 µm from each other, first 3 slices). Data represent mean with SD of 7 organoids. *p < 0.05; Mann-Whitney unpaired t-test. Note that the large majority of organoids is budded at the start of stretch application. **(C)** Distribution of organoids across z slices within BME domes, related to Figure 1C-D (n = 36 organoids from 3 BME domes). **(D)** Axial strain of budded organoids over time during 0.1 Hz cyclic application of 30% stretch, related to Figure 1H. Strain was calculated as the relative change in organoid length (parallel to stretch) during static image acquisition before and during stretch every 30 minutes. Data represent mean with SD of n=5 individual organoids. RM One-way ANOVA revealed no significant differences (p = 0.8537).

**Figure S2.**
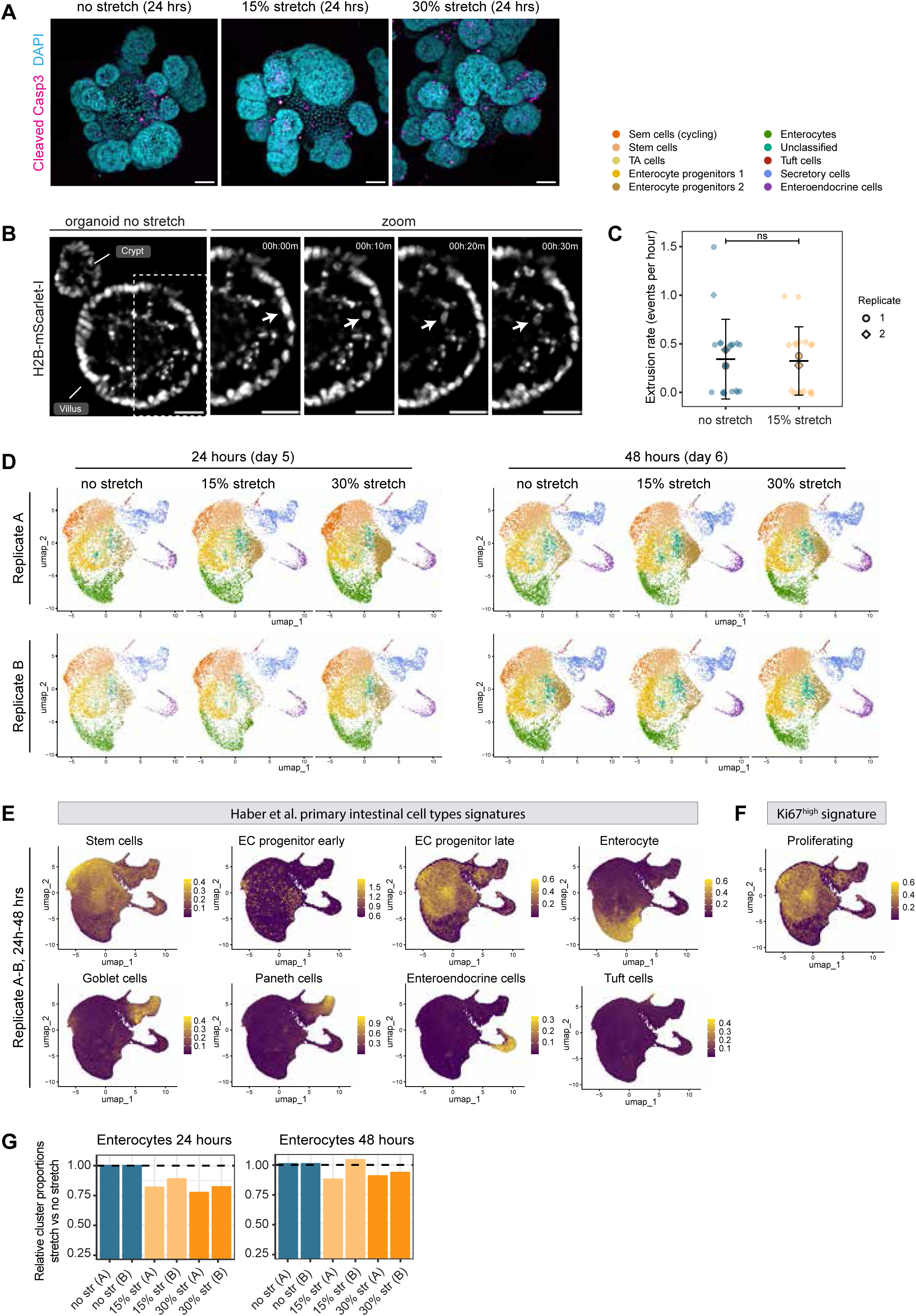
Cell type identification in single cell RNA sequencing data of stretched intestinal organoids. **(A)** Representative maximum intensity z-projection images of intestinal organoids immunostained for Cleaved Caspase-3, together with DAPI, after 24 hours of 15% or 30% cyclic stretch (0.1 Hz), or no stretch. Scale bar: 50 µm. (**B)** Representative timelapse images of an intestinal organoid on PDMS overexpressing H2B-mScarlet-I, with zoom of the villus area showing an extrusion event (white arrow). Time. Scale bar: 30 µm. (**C)** Quantification of the number of extrusion events captured during timelapse imaging of intestinal organoids during 2 hours of 15% cyclic stretching (0.1 Hz) or no stretch, imaged at 1 frame per 10 minutes. Each point represents one organoid (n = 19 without stretch, n = 17 with stretch, from N=2 replicates), with larger points representing replicate means, and horizontal lines overall mean ± SD. Statistics: Poisson generalized linear model with condition and replicate as fixed effects. **(D)** UMAP projections of single cell transcriptomics data from organoids from indicated conditions, showing separate UMAPs for both replicate experiments (A and B). Cluster identities were assigned based on transcriptional similarities to murine intestinal cell type signatures (Haber et al., 2017). **(E)** UMAP projections of pooled single-cell transcriptomes (across conditions, timepoints, and replicates), colored by module scores for gene signatures corresponding to primary murine intestinal cell types (Haber et al., 2017). **(F)** UMAP showing module scores for a proliferation-associated gene signature (derived from Mki67+ cells; Basak et al., 2014) across sequenced cells from all conditions, timepoints and replicates. **(G)** Fold-change proportions of enterocyte cluster cells from 15% and 30% stretched organoids relative to non-stretched organoids at 24 and 48 hours. Bars of the same color represent biological replicates normalized to their corresponding static controls.

**Figure S3.**
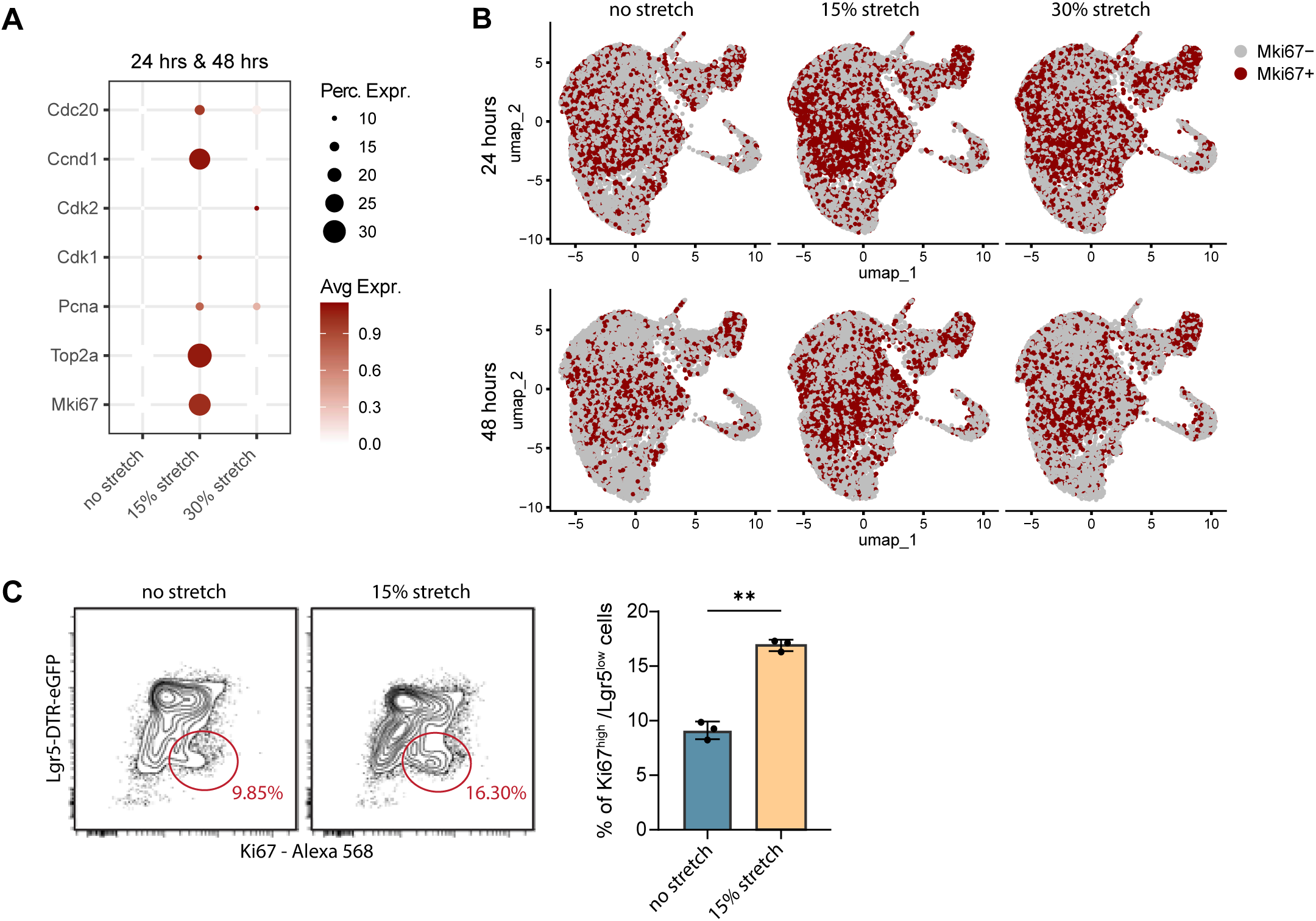
Mechanical strain increases expression of proliferation-associated genes. **(A)** Dot plot showing expression of proliferation markers across single-cell transcriptomes from intestinal organoids cultured without stretch, or with 15% or 30% cyclic stretch (0.1 Hz). Data from two biological replicates (A and B) and two timepoints (24 and 48 hours) were pooled for this analysis. **(B) U**MAP visualization of color-coded Mki67-expressing cells across single-cell transcriptomes from organoids cultured without stretch or under 15% or 30% cyclic stretch (0.1 Hz) for 24 or 48 hours (replicates A and B pooled). Cells with normalized expression of Mki67 above zero are displayed in red, while cells without detectable Mki67 expression are shown in gray. Increased prevalence of Mki67-positive cells in response to stretch suggests strain-dependent increase in proliferation. **(C)** Left: representative contour plots of mKi67 protein staining (x-axis) versus Lgr5-DTR-eGFP (y-axis) intensity of flow cytometry data of fixed single cells isolated from intestinal organoids cultured without or with exposure to 15% cyclic mechanical strain (0.1 Hz) for 48 hours (after 4 days pre-culturing). Encircled cells represent Mki67^high^ proliferating cells with low expression of Lgr5-DTR-eGFP. Right: quantification of the percentage of Ki67^high^/Lgr5-DTR-eGFP^low^ cells by flow cytometry from intestinal organoids with or without 15% cyclic stretch (0.1 Hz). Data from n=3 independent experiments. Paired t-test: ** p < 0.01.

**Figure S4.**
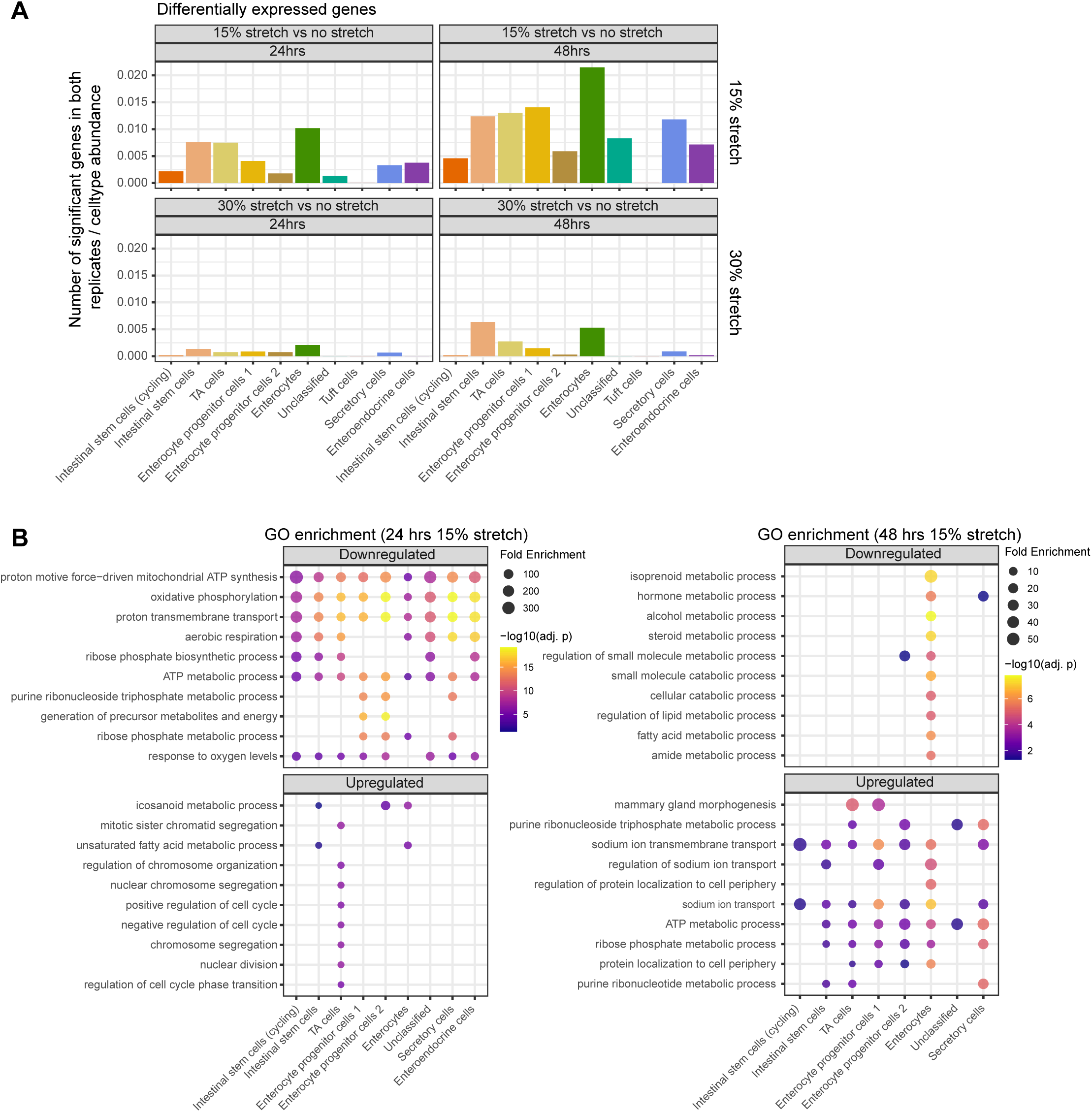
Mechanical strain-induced differentially expressed genes in intestinal organoid cell types. **(A)** Number of differentially expressed genes (pooled up-and downregulated) that were significant in both biological replicate A and B for each cell type cluster, normalized to the abundance of each cell type. **(B)** Top 10 most significantly up-and down-regulated GO terms across all cell types, based on differentially expressed genes between 15% and no stretch after 24 hours (left) and 48 hours (right). Differentially expressed genes were determined using MAST, and only genes significant in both biological replicates were considered.

**Figure S5.**
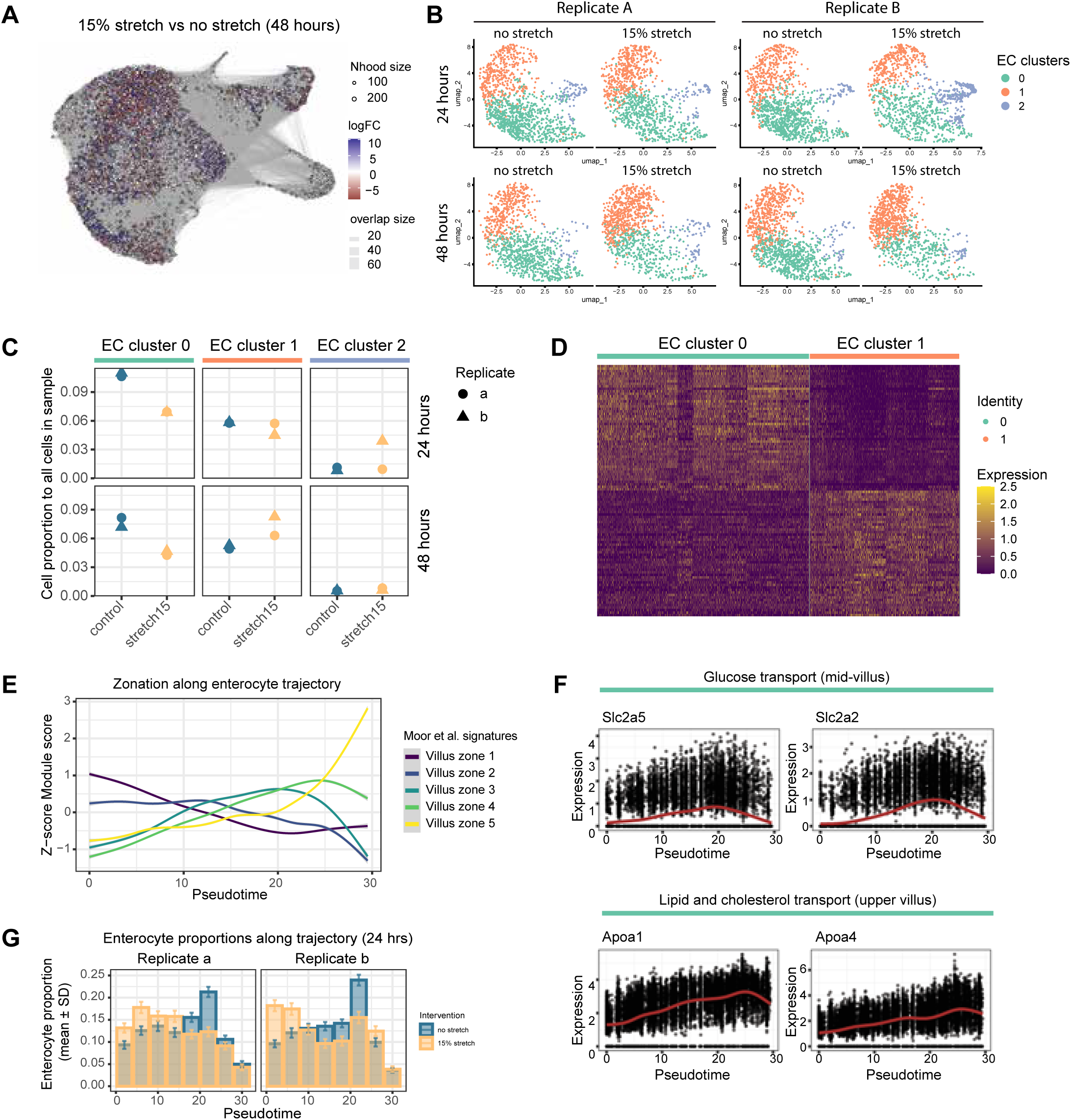
Mechanical strain induces a transcriptional shift from late-to-early stage enterocytes. **(A)** Neighborhood plot showing differential abundances of local neighborhoods plotted on UMAP projection, comparing 48 hours of 15% stretch to no stretch. Both biological replicates are pooled. Neighborhood analysis was performed using MiloR. **(B)** UMAP of isolated enterocytes from organoids with or without 48 hours of 15% cyclic stretch, separated for both biological replicates. Unsupervised clustering identified three subclusters present under both conditions. **(C)** Proportion of cells in each enterocyte subcluster relative to all cells in the respective sample from organoids without stretch or with 15% stretch at 48 hours. Replicate A and B are indicated by symbol shape. **(D)** Heatmap showing top 50 genes enriched in enterocyte subcluster 0 or 1 compared to each other. Combined data from non-stretched and stretched organoids across timepoints and replicates is used to identify marker genes for each cluster. **(E)** Line plots showing module scores of all five villus zonation gene signatures (Moor et al., 2018) along pseudotime values from trajectory analysis of enterocytes (related to figure 3G, conditions, timepoints and replicates combined). Pseudotime (x-axis) represents inferred differentiation progression with module scores (y-axis) reflecting gradual shifts in zonation signatures across this trajectory. **(F)** Line plots showing normalized expression of individual transporter genes involved in glucose transport (villus zone 3) and lipid/cholesterol transport (zone 4-5, Moor et al., 2018) along pseudotime of enterocyte differentiation. **(G)** Relative distribution of enterocytes along pseudotime bins for non-stretched and 15% stretched organoids at 24 hours, related to figure 3G. Mean and SD were calculated using bootstrapping.

**Figures S6.**
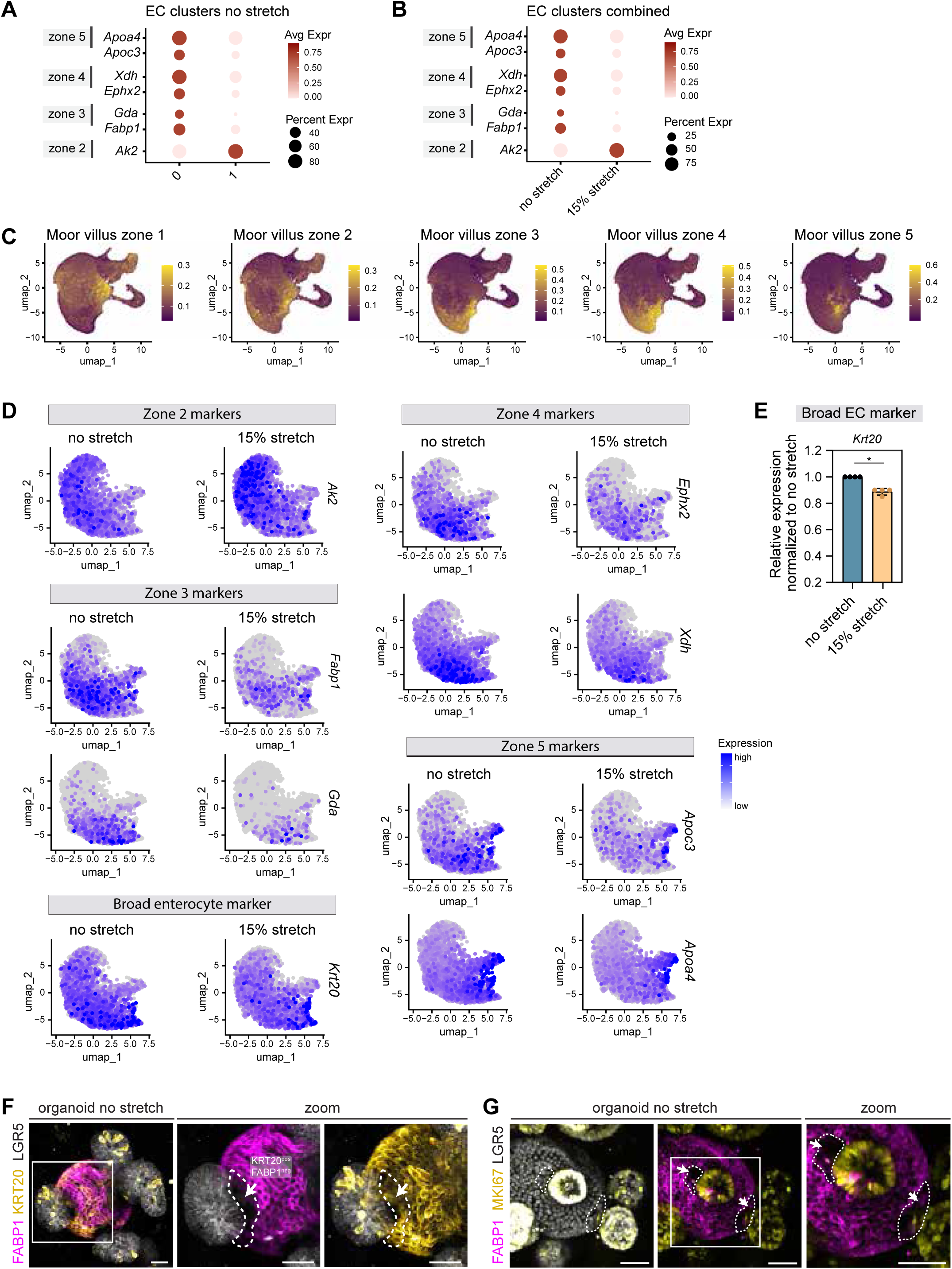
Expression pattern of enterocyte zonation markers across intestinal organoids. **(A)** Dot plot showing expression of zonation marker genes from villus zone 2-5 (Moor et al., 2018) in enterocyte subclusters 0 and 1 from non-stretched organoids (timepoints and replicates pooled). **(B)** Dot plot comparing expression of the same zonation markers (panel A) in all enterocytes (subcluster 0-2 pooled) between non-stretched and 15% stretched organoids (timepoints and replicates combined). **(C)** UMAP projection of colored module scores of villus zonation signatures (Moor et al., 2018) across all single-cell transcriptomes from organoids across conditions, timepoints and replicates. Comparison with figure 2C shows that zone 1 markers are particularly enriched in progenitor cell types outside of the enterocyte cluster. **(D)** UMAPs showing normalized expression of zonation markers from villus zone 2-5 used in RT-qPCR analysis, and broad enterocyte marker Krt20 across enterocytes without versus with 15% stretch (timepoints and replicates pooled). **(E) R**T-qPCR of enterocyte (EC) marker Krt20 in intestinal organoids without stretch or subjected to 15% cyclic stretch for 24 hours. Data are shown as mean ± SD from 4 independent experiments, relative to non-stretched control. Statistics: unpaired Mann-Whitney test on ΔΔCT values per gene; * p < 0.05. **(F)** Representative maximum intensity z-projection image of an intestinal organoid without stretch immunostained for Keratin-20 and mature enterocyte marker FABP1, showing that FABP1-negative cells at the base of the villus (arrow) have started to express general enterocyte marker KRT20. Scale bar: 30 µm. **(G)** Representative maximum intensity z-projection image of intestinal organoid immunostained for proliferation marker MKI67 and mature enterocyte marker FABP1, showing that FABP1-negative enterocytes in the base of the villus are devoid of KI67 (arrows). Scale bar: 50 µm.

**Figures S7.**
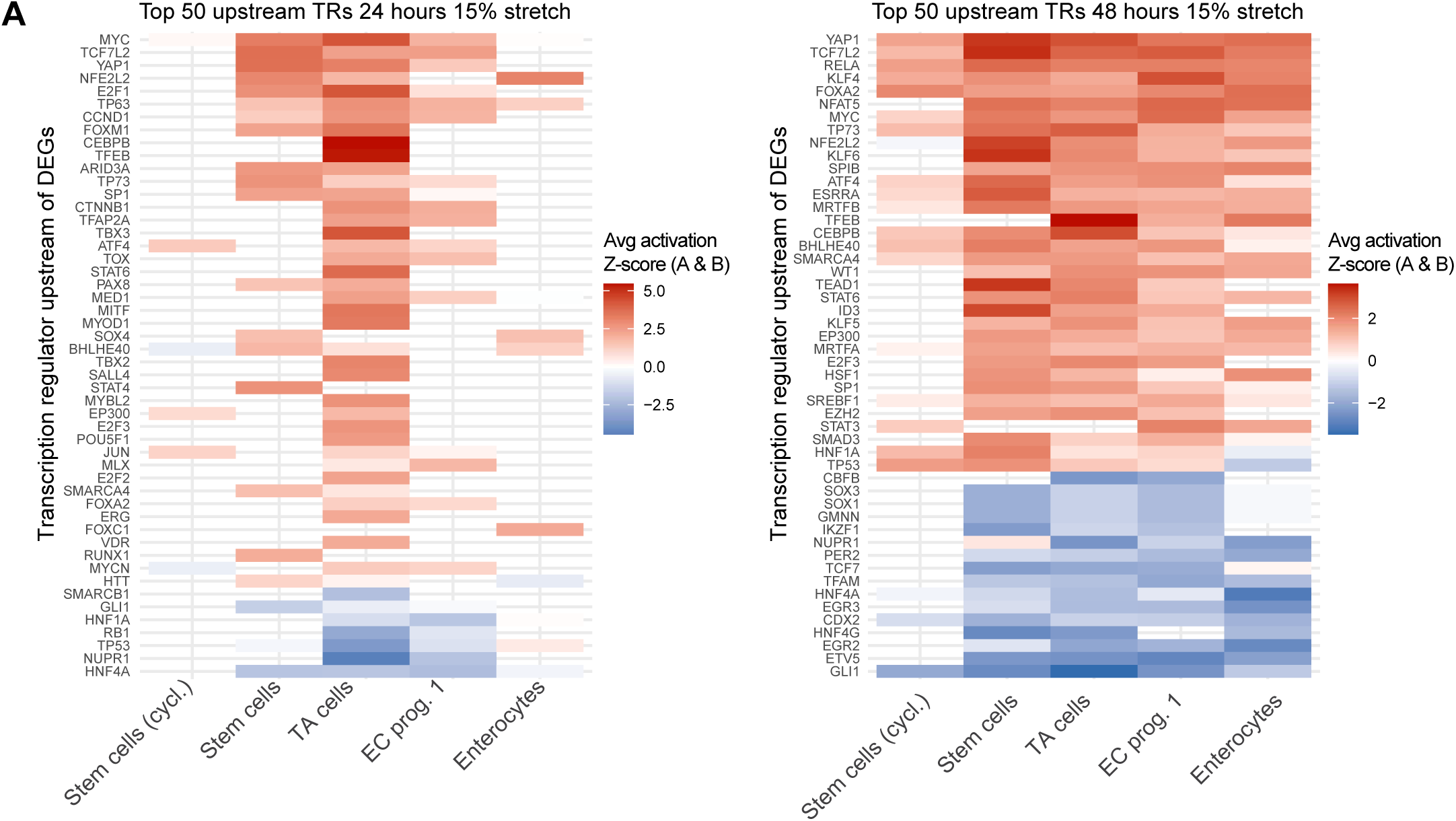
Top 50 identified transcriptional regulators upstream of strain-responsive transcriptomes per cell type. **(A)** Heatmaps displaying average activation Z-scores of upstream regulators across cell types of the absorptive lineage, after 15% stretch for 24 hours (left) and 48 hours (right). Upstream regulators are sorted based on cumulative activation Z-scores across all depicted cell types, and color-coded based on average Z-scores calculated from both biological replicates.

